## Supplementary information for "Comparison between optical tissue clearing methods for detecting administered mesenchymal stromal cells in mouse lungs"

### Supplementary figures

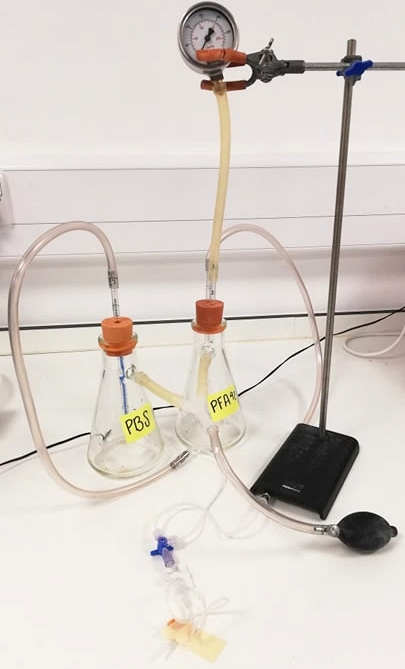

***Supplementary figure 1. Perfusion pump.*** *Two Erlenmeyer flasks, filled with either PBS or 4% PFA, are joined through tubes connected to their sidearms, which are connected to a manual bulb that helps pressurize the system. A pressure gauge is connected* *to one of the bottles to monitor and maintain pressure. Pipettes extend through rubber stoppers and down into the fluid in each flask. The pressure flowing through the sidearm pumps the liquid from each bottle into these pipettes and out of the flasks, where a stopcock allows fluid to flow from one flask at a time.*

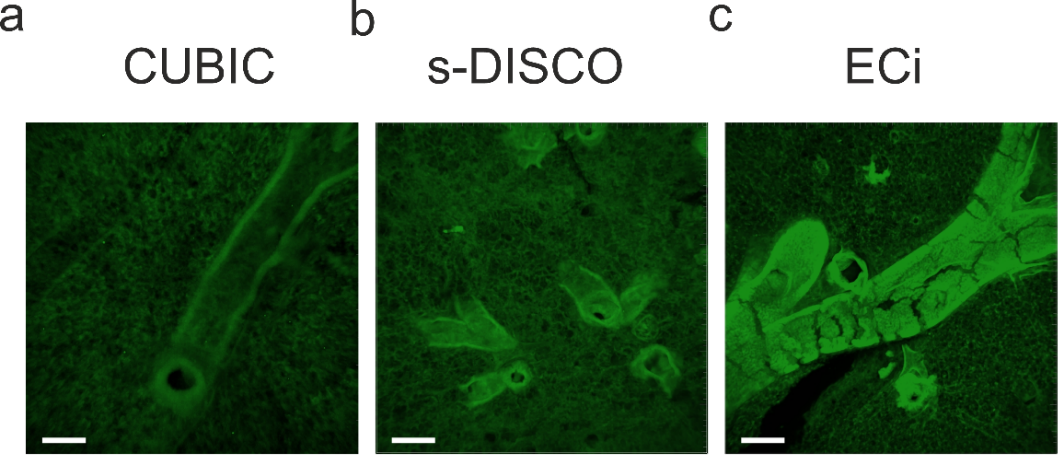

***Supplementary figure 2. Lung morphology after optical tissue clearing.*** *a) CUBIC cleared lung. b) s-DISCO cleared lung. c) ECi cleared lung. Scale bar = 150 µm. MIPs before image reconstruction* *were obtained by confocal imaging on a Leica DMi8 with Andor Dragonfly spinning disk, coupled to an EMCCD camera using a 10x/0.45 air objective. Z-stacks were captured using the 488 nm laser line and 525/50 emission filter.*

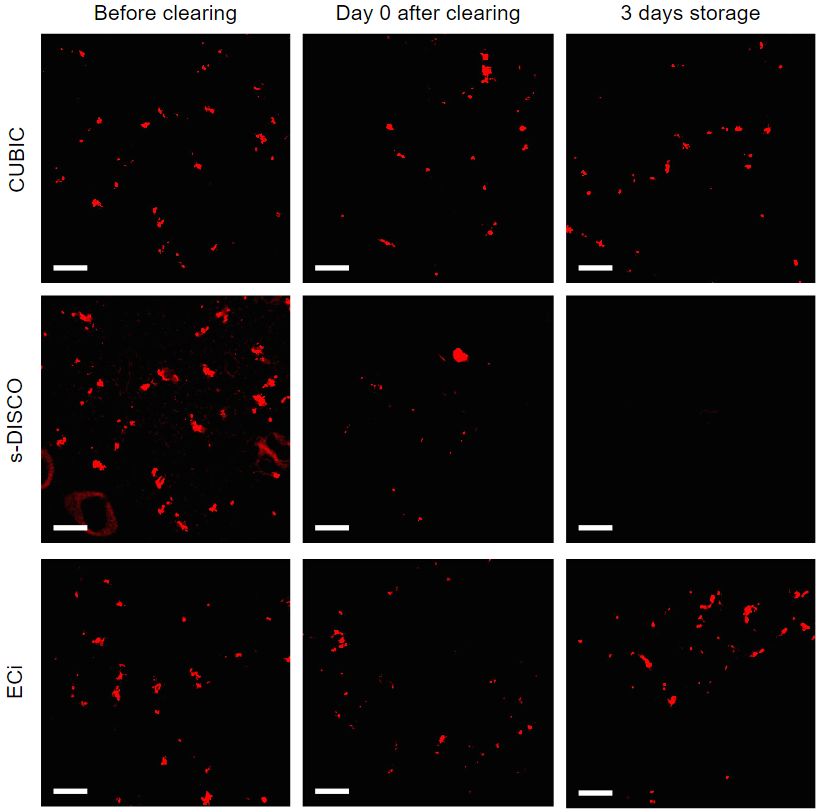

***Supplementary figure 3. Fluorescence preservation of tdTomato (red) after clearing 100 µm thick lung sections.*** *Impact of CUBIC, s-DISCO and ECi clearing on the fluorescence of tdTomato before, immediately after, and 3 days after of storage in RI solution.*

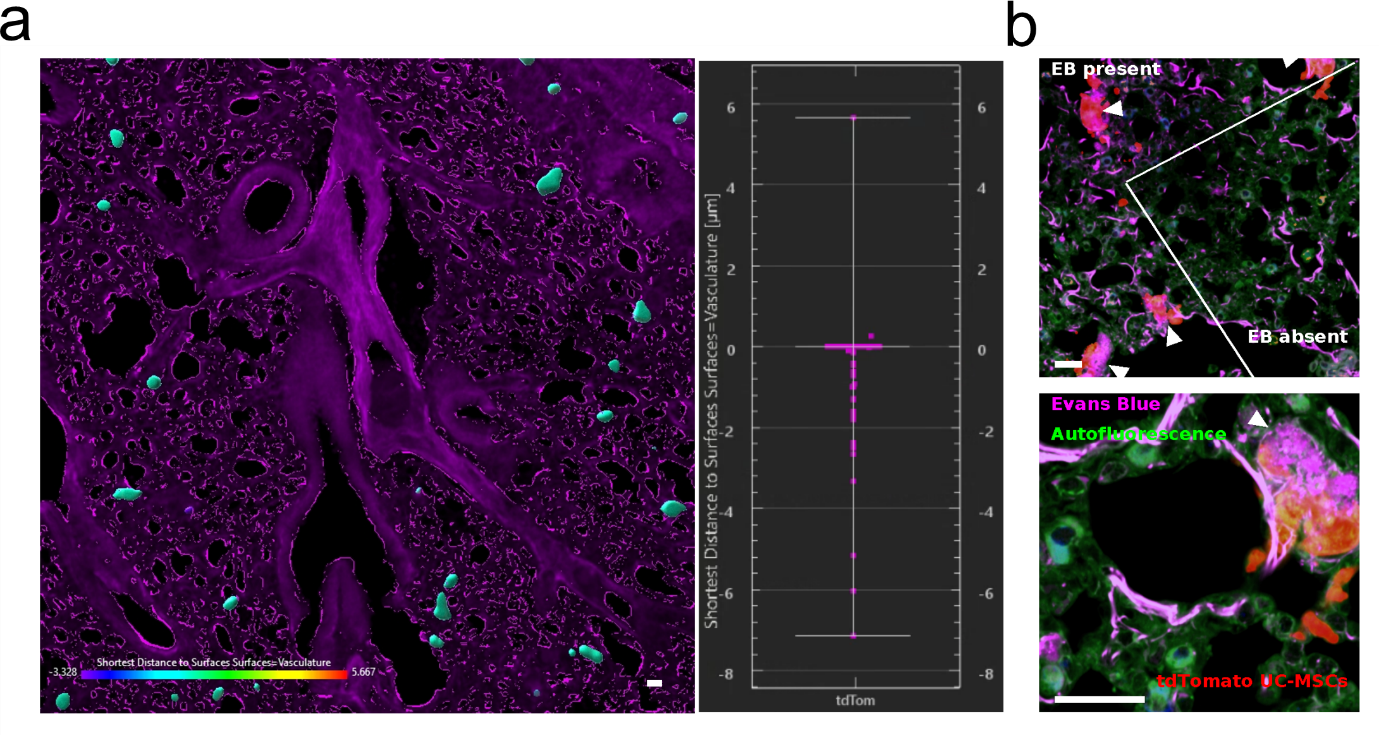

***Supplementary figure 4. Evans Blue vascular staining.*** *a) Lung 30 µm cryosections. IMARIS 3D reconstruction and shortest distance calculation analysis. Distance analysis performed between the tdTomato hUC-MSCs (cyan) and the vasculature (magenta). n = 9. b) tdTomato hUC-MSCs obstructing EB flow through* *vasculature. Arrowheads indicate areas of EB accumulation. Scale bar 25 µm.*

### Supplementary methods

Solvent based optical tissue clearing of 100 µm lung sections.

**Supplementary table 1.** Solvent based optical tissue clearing of 100 µm lung sections.

|  | **100 µm sections** | |
| --- | --- | --- |
|  | **s-DISCO** | **ECi** |
| 1-propanol 50% | 3 min | 3 min |
| 1-propanol 80% | 3 min | 3 min |
| 1-propanol 100% | 3 x 3 min | 3 x 3 min |
| DCM | 1 min | - |
| DBE w/ 0.4% propyl gallate | storage | - |
| ECi | - | storage |
